# An anthropoid/strepsirrhine divergence in ventral visual stream connectivity

**DOI:** 10.1101/2025.05.19.653861

**Authors:** Jasper E. Hunt, Shaun Warrington, Lea Roumazeilles, Saad Jbabdi, Zoltán Molnár, Stamatios N. Sotiropoulos, Rogier B. Mars

## Abstract

The ventral visual stream has undergone extensive reorganisation within the primate lineage. While some work has examined restructuring of the ventral prefrontal cortical grey matter across primates, comparative studies of white matter connectivity are lacking primarily due to difficulties in data acquisition and processing. Here, we present a data-driven approach to the study of white matter connectivity using post-mortem diffusion MRI data. With this approach, we reconstruct anterior temporal-frontal and occipitotemporal-frontal connections across two anthropoids and one strepsirrhine: the rhesus macaque, the black-capped squirrel monkey, and the ring-tailed lemur. We find that the anthropoids exhibit more dorsal prefrontal innervation of these ventral visual connections. This study supports the hypothesis that anthropoid primates underwent extensive reorganisation of both grey and white matter during their emergence as visual foragers in a complex ecological niche. The data-driven techniques presented here enable further research on white matter connectivity in previously understudied species.

## Introduction

The primate brain is a visual brain. Primates are thought to have descended from a common ancestor adapted for life in the fine branches of trees, who developed a leaping-grasping form of locomotion and used frontally directed eyes to guide visual foraging (Cartmill, 1992; Fleagle, 2013; Kaas et al., 2021). This visuomotor history is reflected in the anatomical organisation of the primate brain. One particularly prominent aspect of this is the existence of two cortical pathways in the neocortex. Starting in the striate visual cortex, two visual pathways process different aspects of visual information (Baizer et al., 1991; Goodale & Milner, 1992). The dorsal pathway to posterior parietal cortex processes visual information for visually guided actions (Rizzolatti & Matelli, 2003) while a ventral pathway to inferior temporal cortex was traditionally seen as important in object recognition or more generally the processing of object representations (Kravitz et al., 2013).

Variations in the size of the brain and of aspects of the visual system across the primate order are correlated with the visual specialisations of the different lineages (Barton, 1998). Anthropoid primates in particular moved into the daylight foraging niche, developed a fovea for highly detailed vision, and, in some lineages, trichromatic vision (Williams et al., 2010). As the organisation of an organism’s visual system reflects its ecological niche (Hunt et al., 2024), these anthropoid visual specialisations would be expected to be reflected in the organisation of the cortical visual pathways in anthropoids. However, while the dorsal stream is relatively well studied and its role in the visuomotor niche of primates is well understood, the variation and role of the ventral stream are less often interpreted within such an ecological framework. Recently, however, Murray and colleagues proposed that the role of the ventral stream can be understood in this context as providing the visual context (Murray et al., 2017). They proposed that the inferior temporal cortex provides this information to ventral parts of the prefrontal cortex to guide decision making during foraging and that these connections are particularly elaborated in anthropoids (Eldridge et al., 2021). We undertake a study to provide a first test of this hypothesis.

Understanding the anthropoid elaborations of the general primate neural infrastructure for visuomotor processing requires methods that can study these pathways in a range of species, relatively fast, and at reasonable costs. Detailed tracer studies, although commonly hailed as the gold standard, still face substantial challenges: they require sacrifice of numerous animals per species, are expensive, and are time- and labour-intensive (Nectow & Nestler, 2020). In contrast, recent advances in neuroimaging allow high quality, whole brain data to be collected relatively cheaply either in vivo or from post-mortem brains that are not sacrificed specifically for scientific research (Prescott & Poirier, 2021; Thiebaut de Schotten et al., 2019). Diffusion MRI and related tractography methods in particular have proven successful for comparing brain organisation across species (Hecht et al., 2013; Mars et al., 2018). However, the challenge of such studies is that the brains under investigation are often little studied and have fewer ready-made processing tools available. Standard atlas-based tractography protocols can be used to extract specific pathways, but only if researchers already know a connection’s origin, course, and termination. Such knowledge is increasingly available in human and macaque (Assimopoulos et al., 2024), but by and large absent in other species. As such, without the substantial foreknowledge required for a priori specification of masks and logical operations, a discovery approach is needed to compare brain organisation across species. Exploratory methods for analysing MRI data are well-established (O’Muircheartaigh & Jbabdi, 2018; Singh & Wong, 2010), though their use in comparative studies is still relatively rare (Mars et al., 2019). Data-driven approaches open the unique possibility of interrogating data in lesser-studied species without losing sight of anatomical rigour.

Here, we investigate ventral visual stream connections in the brains of three primate species: the anthropoid Old World macaque monkey, the anthropoid New World squirrel monkey, and the strepsirrhine ring-tailed lemur. We use an unsupervised decomposition algorithm based on non-negative matrix factorization (NMF; Lee & Seung, 1999; Lee & Seung, 2000) to extract connectivity components based on the tractographic inference of white matter trajectories and cortical endpoints (Thompson et al., 2020). We then identify corresponding components between each species, comprising connections between the temporal and prefrontal cortices and connections between the visual and prefrontal cortices. We hypothesise that among these connections, prefrontal innervation is mostly confined to the orbitofrontal cortex in the lemur, but extends more dorsally in the anthropoid primates.

## Methods

### Data

This study used an existing postmortem primate dataset, whose acquisition and early processing has been previously described by Roumazeilles et al. (2022). We used nine post-mortem brains obtained from Copenhagen Zoo, the Zoological Society of London, and the primate research facility at the University of Oxford. These brains comprise three rhesus macaques (*M. mulatta;* 1 female, 2 males; aged 11-15 years); three black-capped squirrel monkeys (*S. boliviensis;* 1 female, 2 males; aged 2-9 years); and three ring-tailed lemurs (*L. catta*, 3 males; aged 3-11 years).

Brains were extracted and formalin fixed. No evidence of neuropathology was found in any of the brains studied. Brains were stored in formalin or a phosphate-buffer saline (PBS) solution. Further details of the brain extraction and storage protocols are reported in Roumazeilles et al. (2022).

### Data preprocessing

Imaging data were collected for all brains using a 7-tesla preclinical MRI scanner (Varian, Oxford UK). All brains were rehydrated in a phosphate buffer for one week prior to scanning, and scanned in Fomblin or Fluorinert. The 7T preclinical scanner used for data acquisition (Varian, Oxford UK) has a birdcage transmit/receive coil with an inner diameter of 72 mm (Rapid Biomedical GmbH, Rimpar Germany).

To minimise image distortions, all 2-D diffusion-weighted images were collected using a spin-echo multi-slice (DW-SEMS) protocol with single line readout. Data are high spatial resolution, at 0.5 mm isotropic resolution for lemurs and squirrel monkeys and 0.6 mm isotropic resolution for macaques, with a single-shell protocol. The acquisition parameters are as follows: TE/TR = 26 ms/10s; matrix size = 128x128 (variable number of slices); b = 0 s/mm^2^ (non-diffusion-weighted; 16 volumes), 4000 s/mm^2^ (diffusion-weighted, 128 volumes). Data quality characteristics including computed signal-to-noise ratio, mean fractional anisotropy, and mean diffusivity are computed and reported for each individual by Roumazeilles et al. (2022).

All data were preprocessed as described by Roumazeilles et al. (2022). Diffusion MRI data were preprocessed using the *phoenix* module of the in-house MR Comparative Anatomy Toolbox (Mr Cat; www.neuroecologylab.org), which relies on FSL (Jenkinson et al., 2012; http://www.fmrib.ox.ac.uk/fsl). A three-fibre ball and stick model was then fitted to the data using FSL’s BedpostX (Behrens et al., 2007). Species-specific brain templates were then created using multimodal iterative registration using the MMORF pipeline (Lange et al., 2020). A cortical surface representation of each template was created using the *precon_all* (Benn et al., 2025; https://github.com/neurabenn/precon_all) adaptation of Freesurfer’s recon-all pipeline (Fischl, 2012) and downsampled to 10,000 vertices per hemisphere.

### Data-driven tractography

We performed data-driven connection mapping using the non-negative matrix factorisation (NMF) algorithm. NMF is an unsupervised dimensionality reduction algorithm with an inherent clustering property, similar to independent component analysis. Briefly, NMF factorises a matrix into the linear product of two matrices, where all three matrices contain only positive values (Figure 1). Because the factor matrices can be lower in dimensionality than the resultant product, NMF generates factors with fewer dimensions than the original matrix. The non-negative nature of NMF makes it well-suited for phenomena which are inherently non-negative (e.g., the firing rates of neurons or the intensity of white matter). In this study, NMF was run via the NFacT pipeline in Python (Thompson et al., 2020; https://github.com/SPMIC-UoN/NFacT), which is a tractography-focused wrapper for Scikit-learn’s (Pedregosa et al., 2011) NMF algorithm.

**Figure 1.**
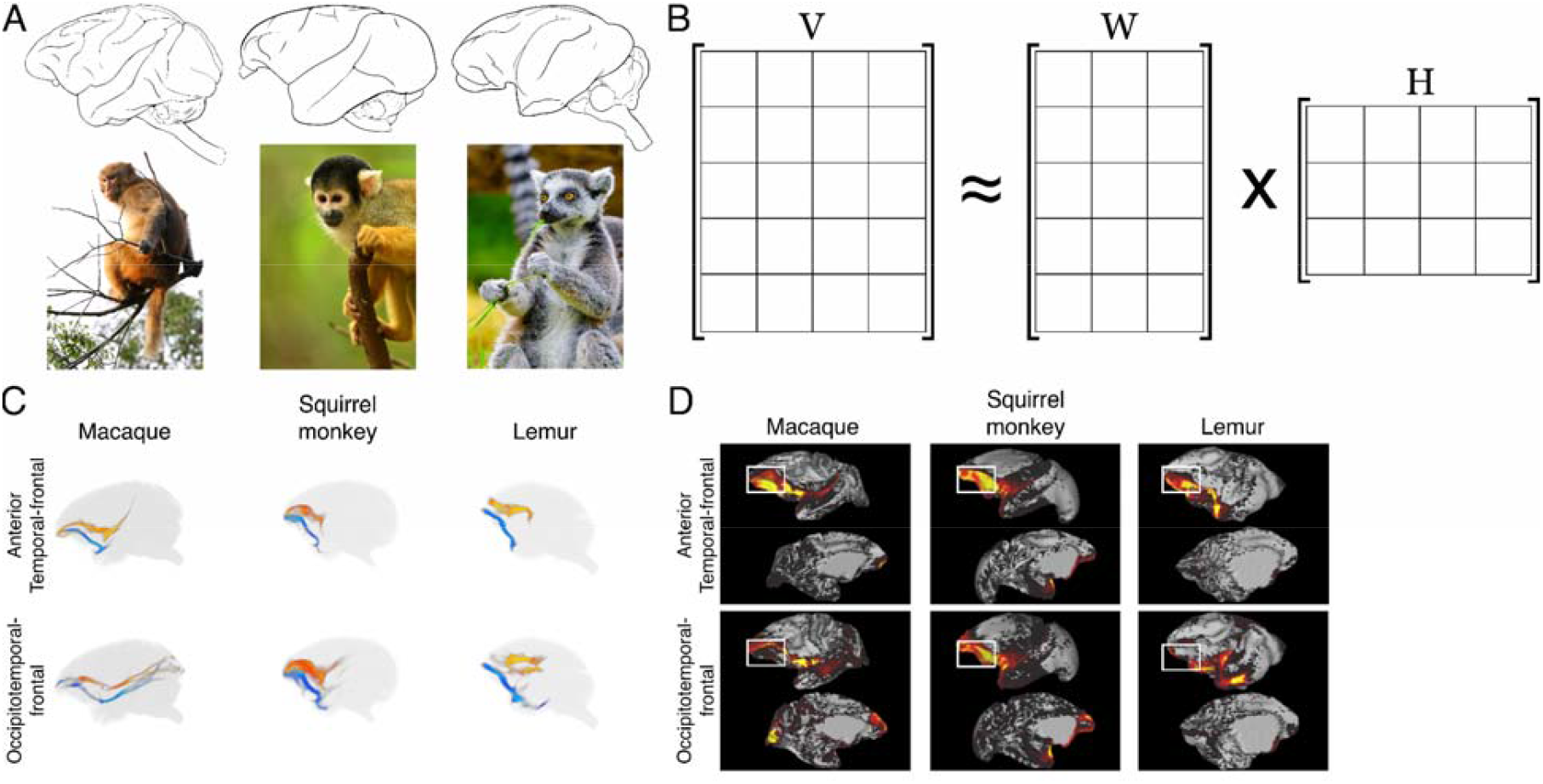
Study overview. **(A)** Diffusion MRI data were collected in three species (left-to-right): the rhesus macaque, the black-capped squirrel monkey, and the ring-tailed lemur. Above: sagittal view of each species’ brain. Below: full-body representation of each species. **(B)** Unsupervised decomposition was conducted to reconstruct white matter connections, using an algorithm called non-negative matrix factorization (NMF). NMF decomposes a matrix, V, into constituent matrices W and H which approximate V when multiplied together. W and H can then be analysed more readily due to their lower dimensionality. **(C)** Ventral visual stream connections with prefrontal termination sites were identified from NMF output in each species. Blue indicates left-hemisphere tracts, while red indicates right-hemisphere tracts. Each 3-D model has been slightly angled from the sagittal plane for visualisation purposes. **(D)** Depth of prefrontal innervation was quantified and compared across species. Each image shows a left-hemisphere brain surface with an example NMF grey matter map overlaid. White boxes illustrate qualitatively that ventral visual stream connections in lemur innervate less dorsal prefrontal cortex than in macaque and squirrel monkey. (Quantitative comparison presented in Results and Figure 4.)

Probabilistic tractography (Behrens et al., 2007) was run from the cortical surface towards the white matter to create a connectivity matrix, wherein each row corresponds to a surface-based (vertex) seed and each column to a volume-based (voxel) target. Individual subject matrices were produced with 10,000 seed points, 1,000 samples per seed, and a step length of 0.2 with distance correction applied, then averaged across individuals. The average connectivity matrix was then decomposed via NMF into two matrices: one comprising white matter by components and one comprising components by grey matter. Thus, for each resultant component, we have both a grey matter surface array of vertices and a volumetric representation of the component’s white matter voxels. These matrices were then regressed from the group decomposition to subject-level decomposition to achieve the individual decomposition data used in analysis.

NMF requires the user to specify the number of components (*K*) to be used in decomposition. Benchmarking work on human diffusion data shows that NMF is able to reconstruct known white matter bundles reliably, but that the number of components in NMF decomposition produces substantial changes to connectivity results (Thompson et al., 2020). Therefore, we perform NMF decomposition for *K* = 50, 60, 100, and 200 components in all brains of our study. To evaluate the performance of data-driven tractography via NMF in the non-human primate, we also performed atlas-based tractography using the macaque monkey brains. We ran probabilistic tractography on a set of 42 combinations of seed, waypoint, and exclusion masks designed to reliably identify the major white matter bundles in the macaque brain, as implemented in the FSL tool *XTRACT* (Warrington et al., 2020). We then ran *XTRACT Blueprint* (Mars et al., 2018; Warrington et al., 2020) to generate surface-space connectivity blueprints. The NMF and XTRACT results were then compared using a Pearson’s correlation, implemented in Python’s numpy library, to generate a correlation matrix.

### Component selection

To reconstruct connections of interest from NMF output, we developed a processing pipeline utilising FSL (Jenkinson et al., 2012), Connectome Workbench (Marcus et al., 2011), and custom tools written in MATLAB (MathWorks, Natick, MA, USA), including the MR Comparative Analysis Toolbox (Mr Cat; Mars et al., 2016; www.neuroecologylab.org).

NMF outputs were analysed to identify and quantify characteristics of the anterior temporal-frontal and occipitotemporal-frontal connections across species. Specifically, regions of interest were drawn on the grey matter surface using Connectome Workbench (Marcus et al., 2011). These regions of interest were specified based on reports in prior literature, summarised in Supplemental Figure 1. In macaque, the orbitofrontal cortex was specified anterior to the infraprincipal dimple in the manner reported by Saleem et al. (2014), who identified the orbitofrontal cortex based on cytoarchitectonic and tracer data in macaques. In squirrel monkey, the orbitofrontal cortex was specified according to Deshpande and Kohut’s (2023) regions of interest, which are based on Yuan and colleagues’ (2021) functional parcellation of the squirrel monkey prefrontal cortex, derived from fMRI data. In lemur, the orbitofrontal cortex was specified as the ventral part of the prefrontal cortex, comprising the grey matter regions ventral to the principal sulcus as specified by Roumazeilles et al. (2022), and bounded posteriorly by the edge of prefrontal cortex that was identified cytoarchitectonically by Mott and Kelley (1908). Temporal regions of interest were drawn to include either the anterior temporal pole or a larger swathe of temporal cortex (see *Supplemental Methods*). Occipital regions of interest used either the primary visual cortical regions of interest reported by Roumazeilles et al. (2022) or larger regions of interest comprising most of the occipital lobe (see *Supplemental Methods*). It was found that the specification of temporal and occipital regions of interest did not affect the selection of anterior temporal-frontal or occipitotemporal-frontal components. The temporal and occipital regions of interest were not used in the calculation of prefrontal innervation metrics or the statistical analysis of cross-species differences.

Using the regions of interest described above, we identified components comprising anterior temporal-frontal and occipitotemporal-frontal connections based on the signal intensity in temporal and prefrontal or occipital and prefrontal surface regions of interest. To identify which components had high signal intensity in both regions of interest, the average signal intensity in each region was computed for each component, and these averages were multiplied to produce plots showing which NMF components have high signal intensity in both regions of interest. Those components which had high signal intensity in both regions were then visualised in both surface and volume space, to identify anterior temporal-frontal and occipitotemporal-frontal connections. Because the connections we studied are all ipsilateral, contra-hemispheric values were zeroed at this point to aid in visualisation. Identification of connections took place on the basis of the following criteria. Anterior temporal-frontal connections were identified based on termination sites in the anterior temporal pole and prefrontal cortex, a hook-shaped morphology around the lateral sulcus, minimal parietal involvement, and minimal occipital involvement. Occipitotemporal-frontal connections were identified based on termination sites in the ventral occipital cortex and prefrontal cortex, a morphology running principally along the anterior-posterior axis, minimal parietal involvement, and a ventral position in volume space. It has been previously reported that at higher NMF component numbers, fibre bundles split into multiple sub-components that together comprise a single projection of interest (Thompson et al., 2020). We found this to be the case as well. To counteract this issue, whenever multiple components matched the anatomical criteria outlined above at higher NMF model orders, these components were summed to produce a single connection.

### Statistical analysis

After identifying anterior temporal-frontal and occipitotemporal-frontal connections for each species and each number of components, we quantified the distribution of termination sites in the prefrontal cortex, enabling comparison of the depth of prefrontal innervation across species. For each brain, an orbitofrontal bias index (OBI) was computed. This index is defined by dividing the signal intensity of a connection’s termination sites in the orbitofrontal cortex by the signal intensity of all of that connection’s termination sites in the prefrontal cortex. The resultant value is divided by the relative size of the orbitofrontal cortical region of interest compared to the rest of prefrontal cortex. In other words, OBI is the percentage of a connection’s prefrontal termination which lies within the orbitofrontal cortex, normalised to the relative size of the orbitofrontal cortex. A high OBI value means the reconstructed connection terminates largely in the orbitofrontal cortex. A low OBI value means relatively more of the connection terminates dorsal to the orbitofrontal cortex.

To compare OBI values, a nested ANOVA was implemented using the spreadsheet provided by the *Handbook of Biological Statistics* (3rd ed.) (McDonald John H., 2014). This approach allows for statistical comparison of the contribution to variance by cross-species differences (macaque, squirrel monkey, or lemur); within-species differences; and NMF *K* value. This comparison yields a percentage of variance explained by each hierarchical variable in the nested ANOVA. In other words, nested ANOVA allows us to identify what percentage of variation in our dataset comes from cross-species differences, what percentage originates from differences between brain hemispheres, and what percentage originates from adjusting NMF component numbers. To compute statistically significant differences between groups, post-hoc analyses were completed by t-test with a Bonferroni correction for multiple comparisons.

## Results

In this study, we identify and characterise differences in the prefrontal innervation of anterior temporal-frontal and occipitotemporal-frontal connections between two anthropoid species and one strepsirrhine species: the rhesus macaque, the black-capped squirrel monkey, and the ring-tailed lemur (Figure 1A). These connections are reconstructed using a data-driven method based on non-negative matrix factorisation (NMF; Figure 1B).

First, we benchmark the performance of NMF against traditional tractography methods. We then identify corresponding anterior temporal-frontal and occipitotemporal-frontal connections in each species (Figure 1C). Next, we quantify and compare the prefrontal innervation of both connections in each species (Figure 1D). Finally, we assess the contributions to overall variance in the data from cross-species differences, within-species differences, and algorithmic parameterisation, to conclude that the cross-species differences reported here are far greater than individual (within-species) differences or methodological-based differences.

### NMF output closely recapitulates known white matter tracts in macaque

NMF can be used to perform data-driven tractography which closely recapitulates known white matter tracts in human (Thompson et al., 2020). Here, we extend this finding to macaque, demonstrating that even with small sample sizes, it is possible to reproduce known white matter tracts in a non-human primate for which a *priori* described tractography protocols also exist (Assimopoulos et al., 2024; Mars et al., 2018; Warrington et al., 2020).

Because NMF and related algorithms require users to specify a number of components for parcellation, we ran NMF with a range of component values, from *K* = 10 to *K* = 200 components (i.e., parcellating the subject’s brain into 10 to 200 components), using the NFacT toolbox (https://github.com/SPMIC-UoN/NFacT). Each NMF component comprises a white matter volumetric map and a grey matter endpoint map, representing grey matter regions connected by the bundles depicted in the white matter map. At all component numbers tested, we were able to identify components which recapitulate well-known tracts in both white matter and grey matter properties, including the uncinate fasciculus and the inferior fronto-occipital fasciculus (Figure 2). For K values ranging from 10 through 200, we identified the best-correlated NMF component with each of the 42 white matter tracts identified using XTRACT’s tractography protocols for macaque (Warrington et al., 2020). These correlations are presented in Figure 2A. Consistent with previous findings (Thompson et al., 2020), we observe that at low component numbers, components tend to include white matter that is nonspecific to the tracts of interest and at high component numbers, components are more likely to correspond to subdivisions of tracts (Figure 2B).

**Figure 2.**
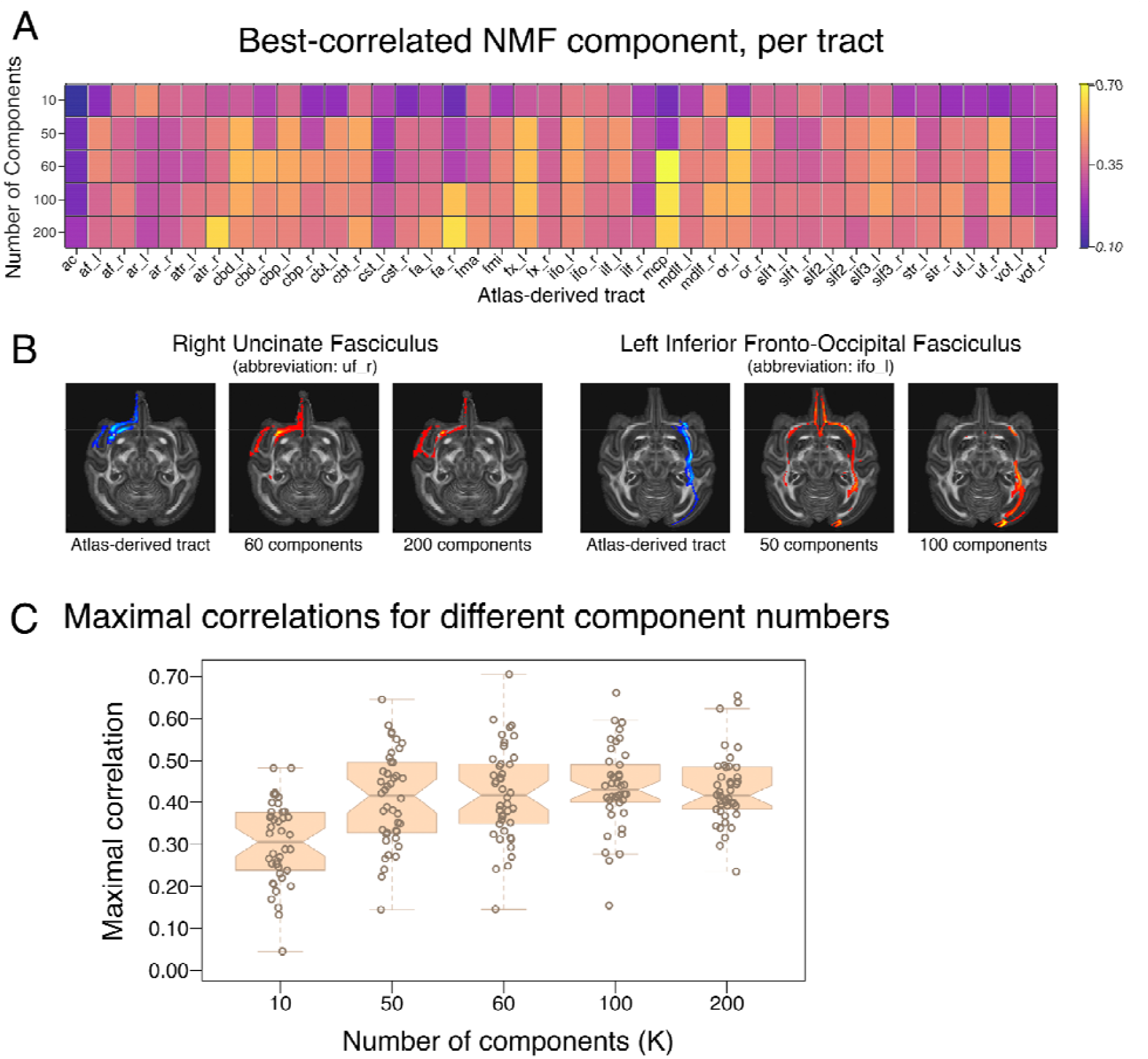
Non-negative matrix factorisation (NMF) can reconstruct known white matter tracts in macaque. **(A)** Table of maximal correlations between NMF components and atlas-derived tracts. Each cell represents the correlation of the best-matching NMF component for a given atlas-derived tract (columns), at different numbers of components (rows). Abbreviations correspond to FSL naming conventions (Jenkinson et al., 2012) and XTRACT output file structures (Warrington et al., 2020; https://fsl.fmrib.ox.ac.uk/fsl/fslwiki/XTRACT). (*Figure 2, cont*.) **(B)** Representative examples of ventral visual stream tracts reconstructed from NMF components. At lower component numbers, NMF output tends to include spurious white matter which does not comprise the projection of interest, while at higher component numbers, NMF output tends to split fibre bundles into multiple sub-components that together comprise a single projection of interest. Blue: atlas-derived tract. Red: NMF-derived tract. **(C)** Distributions of maximal correlations for different numbers of NMF components. Box-and-whisker plots show the distributions of maximal correlations for different numbers of components. Box-and-whisker plot notches indicate the 95% confidence interval of the median (median ± 1.57 * IQR/n^0.5^).

Having established that NMF can reliably reproduce known white matter tracts in macaque, we characterised the parameters under which NMF is most suitable for atlas-based tractography in non-human brains. Other authors (Thompson et al., 2020; Varikuti et al., 2018) have reported that NMF tends to output components which better mirror brain features of interest when run with a number of components that closely matches the number of features desired. When *K* is substantially less than or greater than the number of connections to be reconstructed, the best-matching NMF components tend to be less well correlated with the ground-truth tracts in human subjects (Thompson et al., 2020). Qualitatively, our data paint a similar picture in macaque (Figure 2B). Although maximal correlation distributions were similar for all *K* values ranging from 50 through 200 (Figure 2C), the best-suited value for *K* varied on a per-tract basis (Figure 2A); this complexity indicates that the most appropriate number of components for white matter parcellation could vary from tract to tract, species to species, and brain to brain. Indeed, in applying the related algorithm, orthogonal projective nonnegative matrix factorization, to parcellate grey matter across species, (Vickery et al., 2024) report that the most appropriate number of components differed between species. To guard against the possibility of error or bias arising from number of components specified in the NMF algorithm, all subsequent experiments reported here performed NMF decomposition with 50, 60, 100, and 200 components and the results were compared (see section, *Cross-species differences explain far more variance than within-species differences or algorithmic parameters*).

### Anterior temporal-frontal and occipitotemporal-frontal connections can be reliably reconstructed across primate species

To reconstruct ventral visual stream connections from NMF output, we developed the following processing pipeline. First, NMF output was sorted to find components comprising the connections of interest. A custom MATLAB script was run to identify which NMF components have grey matter maps with high signal intensity in specific regions of interest (Figure 3A-B). These regions of interest were identified based on anatomical landmarks and previous literature (*see Methods*, Supplemental Figure 1). Specifically, anterior temporal-frontal connections were screened based on high signal intensity in the anterior temporal pole and the prefrontal cortex, while occipitotemporal-frontal connections were screened based on high signal intensity in primary visual cortex and prefrontal cortex. It should be noted that an additional analysis in which these regions of interest were expanded to include more occipital and temporal cortex did not yield any additional components comprising the connections of interest (*see Supplemental Methods*).

**Figure 3.**
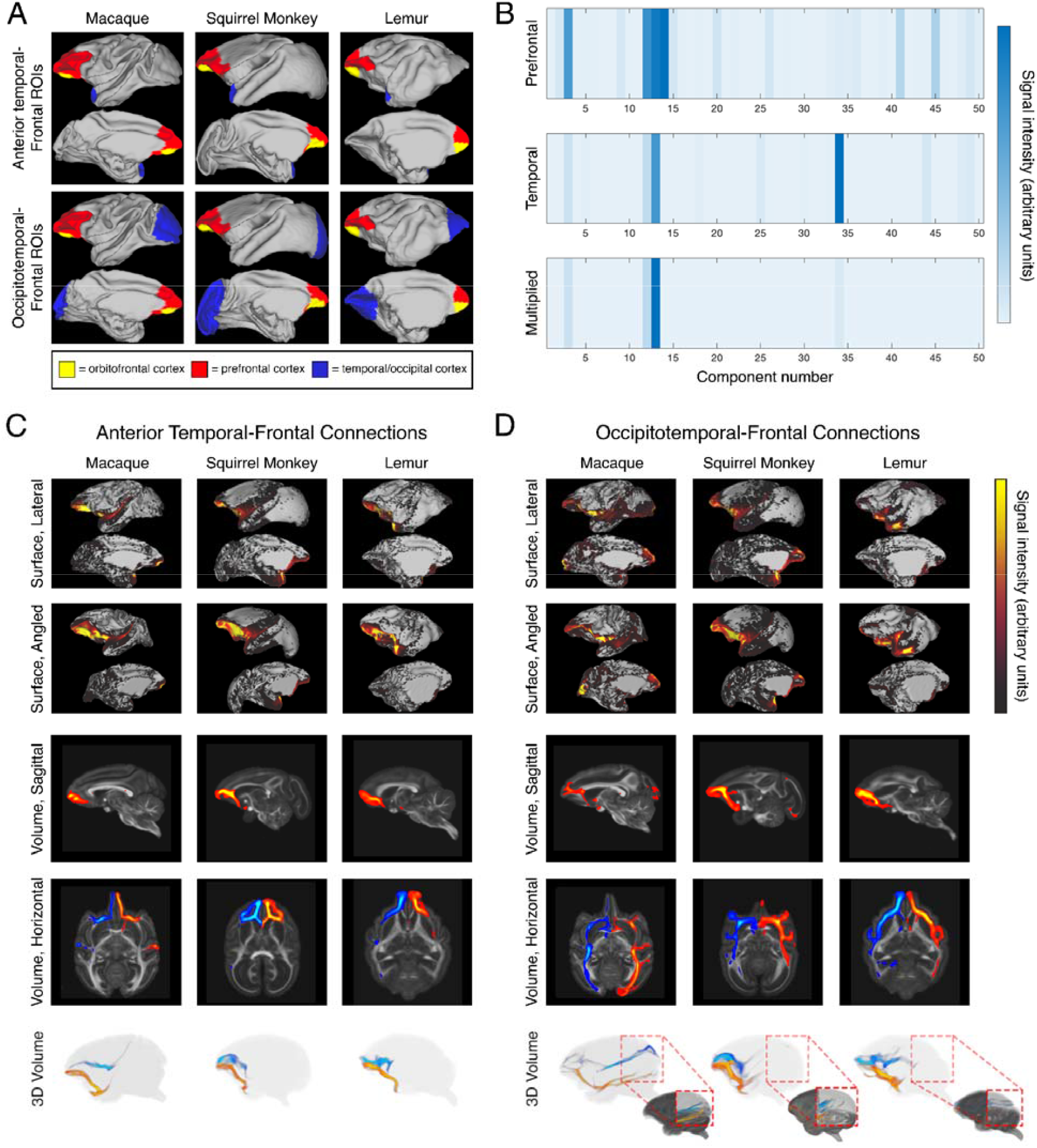
Reconstruction of anterior temporal-frontal and occipitotemporal-frontal connections. **(A)** Surface projections showing regions of interest used to screen NMF (*Figure 3, cont*.) components. Yellow: orbitofrontal cortex; red: prefrontal cortex; blue: anterior temporal pole (top), primary visual cortex (bottom). Note: the orbitofrontal cortex is a subregion of the prefrontal cortex; as such, all analyses of the prefrontal cortex include both the yellow and red regions. **(B)** Sample output of surface intensity screening. Top: each component’s intensity in the prefrontal region. Middle: each component’s intensity in the temporal region. Bottom: intensity values from each region multiplied. Components with high intensity values in both regions were selected for visualisation to identify putative connections. **(C)** Example anterior temporal-frontal connections in each species. Top two rows: surface intensity projections showing putative innervation of reconstructed connections. Both rows present the lateral surface (above) and medial wall (below) of the left hemisphere. The first row is a sagittal view. The second row is the same surface, with the orientation slightly tilted to show more of the prefrontal surface. Bottom three rows: volumetric reconstructions of anterior temporal-frontal connections in a left hemispheric sagittal plane (row three); horizontal plane in radiological orientation (row four); and 3D space (row five). The 3D volumetric reconstruction has been slightly angled from the sagittal plane, to better visualise the bilateral morphology of reconstructed connections. For bilateral reconstructions in rows four and five, blue indicates a right hemispheric connection while orange indicates a left hemispheric connection. **(D)** Example occipitotemporal-frontal connections in each species. Rows are as described in panel C. Row five includes a threshold-adjusted inset volume, to visualise the posterior part of each connection.

Following sorting, components with high signal intensities in the relevant grey matter cortical regions were visually inspected in both surface and volume space to identify those that comprise anterior temporal-frontal and occipitotemporal-frontal connections. Anterior temporal-frontal connections were identified based on the following anatomical criteria: minimal parietal involvement, minimal occipital involvement, termination sites in the anterior temporal pole and prefrontal cortex, and a hook-shaped morphology around the lateral sulcus.

Occipitotemporal-frontal connections were identified based on the following criteria: minimal parietal involvement, termination sites in the ventral occipital cortex and prefrontal cortex, and a morphology running principally along the anterior-posterior axis. Where connections had been split into multiple components – primarily at high NMF component numbers – these components were then merged to form the complete connection. In each species, the reconstructed anterior temporal-frontal connections bore a strong resemblance to a known tract in the uncinate fasciculus (Figure 3C). Occipitotemporal-frontal connections resembled the inferior fronto-occipital fasciculus, but showed more considerable variation in constitution and may have involved other occipitotemporal-frontal connections (Figure 3D). Each connection is additionally presented as a series of sections across coronal, axial, and sagittal planes in Supplemental Movies 1-9.

### Anterior temporal-frontal connections innervate more dorsal prefrontal cortex in anthropoid primates

Having reconstructed anterior temporal-frontal connections in each species, we quantified the depth of prefrontal innervation in each species using a novel measure called the orbitofrontal bias index (OBI). OBI is a measure of the tendency for a connection to terminate within the orbitofrontal cortical region on the ventral side of the prefrontal cortex. Specifically, OBI is defined as the proportion of signal intensity in the orbitofrontal cortex compared to the rest of prefrontal cortex, divided by the size of the orbitofrontal cortex compared with the rest of prefrontal cortex. A high OBI value means the anterior temporal-frontal connection terminates largely in the orbitofrontal region. A low OBI value means relatively more of the anterior temporal-frontal connection terminates dorsal to the orbitofrontal region. In this way, the OBI allows for quantitative comparison of how a connection innervates the different subdivisions of the prefrontal cortex.

After calculating an OBI value for each reconstructed anterior temporal-frontal connection, these data were arranged in a hierarchical fashion, with top-level groups comprising different species and sub-groups comprising individual brain hemispheres (Figure 4A). Because NMF was run on each brain a total of four times at different component numbers, this hierarchical ordering of data points was chosen to respect the assumption of statistical independence among data points during inferential statistical tests. During comparisons between brain hemispheres or between species, the individual data points that arise from repeated sampling are averaged together. (See Methods for further detail.)

**Figure 4.**
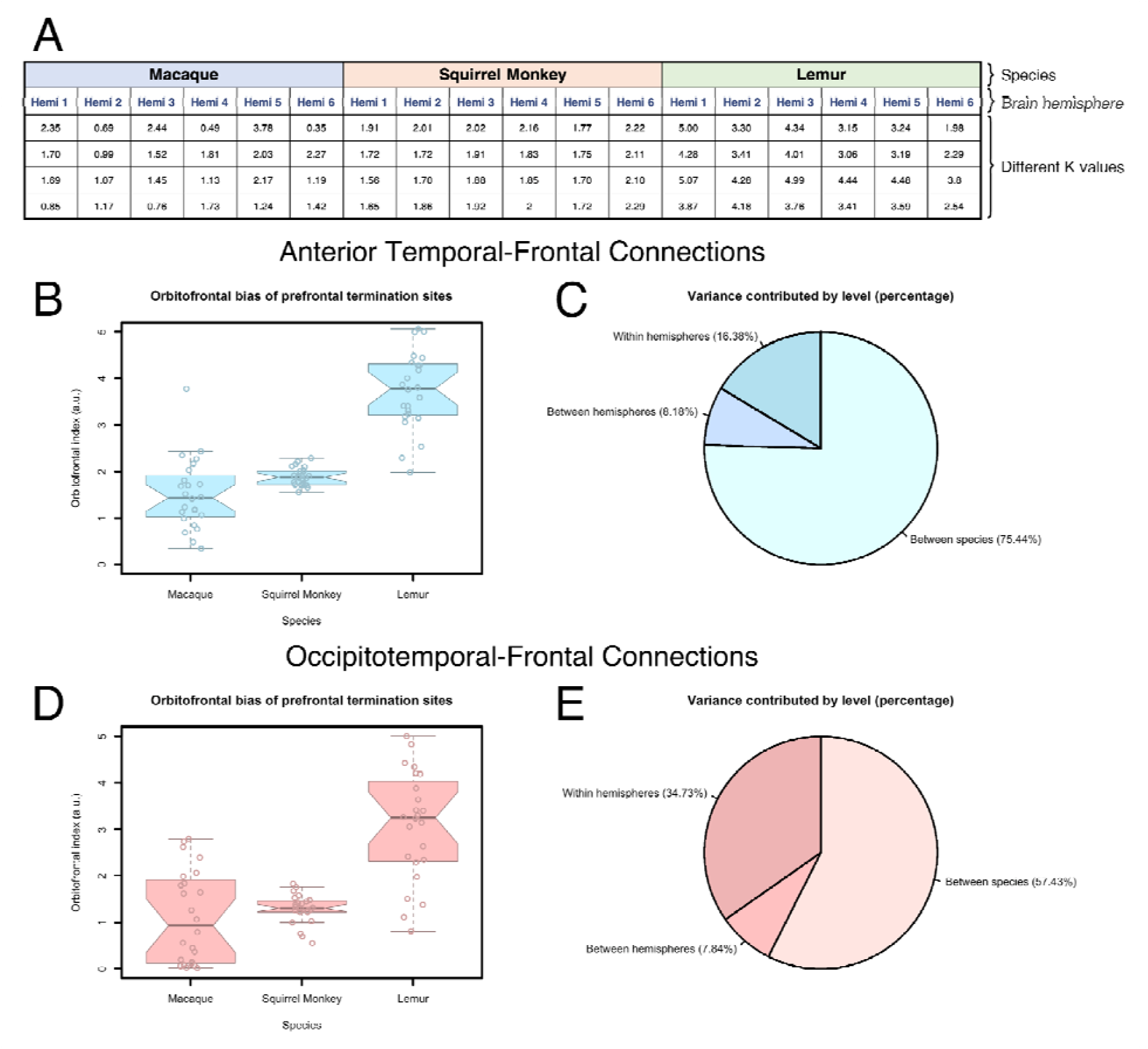
Lemur shows the highest orbitofrontal bias in both sets of connections, with cross-species differences being far larger than other sources of variance. **(A)** Hierarchical organisation of the nested ANOVA. The top level comprises different species: macaque, squirrel monkey, and lemur. The middle level comprises groupings by individual brain hemisphere. The (*Figure 4, cont*.) lowest level comprises individual data points derived from running NMF with different numbers of components. **(B)** Anterior temporal-frontal orbitofrontal bias index values differ in lemur from macaque and squirrel monkey. Box-and-whisker plots show the distributions of orbitofrontal bias index values. Notches indicate the 95% confidence interval of the median (median ± 1.57 * IQR/n^0.5^). **(C)** Different contributions to variance in anterior temporal-frontal orbitofrontal bias index values. Between-species differences comprise 75.44% of the variance in the dataset, while between-hemisphere differences comprise 8.18% and NMF parameterisation comprises 16.38%. **(D)** Occipitotemporal-frontal orbitofrontal bias index values differ in lemur from macaque and squirrel monkey. Box-and-whisker plots show the distributions of orbitofrontal bias index values. Notches indicate the 95% confidence interval of the median (calculated as in Panel B). **(E)** Different contributions to variance in occipitotemporal-frontal orbitofrontal bias index values. Between-species differences comprise 57.43% of the variance in the dataset, while between-hemisphere differences comprise 7.84% and NMF parameterisation comprises 34.73%.

Anterior temporal-frontal connections in macaque and squirrel monkey had similar OBI values (mean ± standard deviation: 1.51 ± 0.76 for macaque; 1.89 ± 0.19 for squirrel monkey). In contrast, lemur had an anterior temporal-frontal OBI value of 3.74 ± 0.82 (n = 24 observations from 6 brain hemispheres per species; distributions presented graphically in Figure 4B). Using a nested ANOVA for inferential statistics, we observed a significant difference between species on anterior temporal-frontal OBI values (*F*: 37.87, *d*.*f*. = 2, *p* = 1.37 * 10^-6^). Post-hoc t-tests with the Bonferroni correction showed that macaque and squirrel monkey differed from lemur (*p* = 2.81 * 10^-12^ and p = 1.39 * 10^-13^, respectively), but not from each other (*p* = 0.07).

One possible confound in this analysis is the specification of orbitofrontal regions. Because the ventrolateral prefrontal cortex expanded in anthropoids **(Eldridge et al**., **2021)**, the orbitofrontal cortex constitutes a relatively smaller portion of the prefrontal cortex in these species. The size of our orbitofrontal regions of interest reflects this, with macaque’s being the smallest relative to the size of the full prefrontal cortex. Although the OBI measure controls for this by dividing the relative signal by the relative size of the orbitofrontal cortex, it is possible that defining a smaller region of interest in macaque could lead to artificially deflated OBI values. As an additional control against this possible confound, the same analysis was run with a more liberally-defined orbitofrontal region of interest in macaque, and the resulting OBI values analysed independently from the previous results (see Supplemental Methods, Supplemental Figure 2). The results of both analyses agree, with lemur once again having the highest average OBI value by a sizable margin (lemur: 3.74 ± 0.82, compared with 2.61 ± 0.62 for macaque and 1.89 ± 0.19 for squirrel monkey). In this additional control experiment, Bonferroni post-hoc revealed significant differences between all three species (macaque v. lemur: *p* = 8.15 * 10^-6^, squirrel monkey v. lemur: *p* = 1.39 * 10^-13^, macaque v. squirrel monkey: 6.25 * 10^-6^).

### Cross-species differences explain far more variance in anterior temporal-frontal connectivity than within-species differences or algorithmic parameters

Using a nested ANOVA design, we quantified how different factors contribute to overall variance within our dataset. Because of the hierarchical organisation of a nested ANOVA, it is possible to make statistical comparisons of the contribution to variance by each level in the hierarchy. By organising our nested ANOVA first by species (macaque, squirrel monkey, or ring-tailed lemur), then different brain hemispheres, then NMF component number, our analysis yields a percentage of variance explained by each hierarchical level in the nested ANOVA. In other words, the nested ANOVA allows us to identify what percentage of variation in our dataset comes from cross-species differences, what percentage originates from differences between brain hemispheres of the same species (e.g., due to age, sex, or hemispheric asymmetries), and what percentage originates from adjusting NMF component numbers.

Differences between species accounted for 75.44% of the variance in the dataset, whereas within-species differences accounted for only 8.18% of the variance. Running NMF with different numbers of components contributed the remaining 16.38% of the variance (Figure 4C). By running NMF with different numbers of components and assessing the contribution of different algorithmic parameters to overall variance in the dataset, we can conclude that the cross-species differences observed here are several times larger than inter-individual differences, inter-hemispheric differences, and variability in the identification of connections arising from the use of different NMF parameters.

### Occipitotemporal-frontal connections innervate more dorsal prefrontal cortex in anthropoid primates

Our anterior temporal-frontal results are consistent with previous work suggesting that anthropoid primates evolved to rely more on ventrolateral prefrontal areas for visual processing in the ventral visual stream. However, it is uncertain whether other ventral visual processing tracts follow the same pattern. To address this question, we quantified the prefrontal innervation of occipitotemporal-frontal connections in the same way as we did anterior temporal-frontal connections.

We generated orbitofrontal bias index (OBI) values for each species’ reconstructed occipitotemporal-frontal connections following the same procedure as we did for anterior temporal-frontal connections. In every species, OBI values were lower for the occipitotemporal-frontal connections than they were for anterior temporal-frontal connections. Again, macaque and squirrel monkey showed similar OBI values (mean ± standard deviation: 1.17 ± 0.99 for macaque; 1.29 ± 0.31 for squirrel monkey), while OBI values in lemur were substantially higher at 3.08 ± 1.17 (n = 24 observations from 6 brain hemispheres per species; distributions presented graphically in Figure 4D). A nested ANOVA revealed a significant difference between species, albeit with a lower F-statistic than for anterior temporal-frontal analysis (*F*: 21.86, d.f. = 2, *p* = 3.6 * 10^-5^), meaning that between-species differences explain less of the overall variance in the dataset. Post-hoc *t*-tests with Bonferroni correction showed that differences between lemur and the two anthropoid species were the principal driver of the ANOVA’s overall effect (macaque v. lemur: *p* = 3.28 * 10^-7^; squirrel monkey v. lemur: *p* = 1.23 * 10^-8^; macaque v. squirrel monkey: *p* = 1.00).

Consistent with the lower *F*-statistic, between-species differences accounted for a relatively lower proportion of the overall variance in the dataset at 57.43%, while within-species differences contributed 7.84% of the variance and specifying different *K* values for NMF contributed 34.73% of the variance (Figure 4E). In sum, these results reflect that lemur occipitotemporal-frontal connections are substantially more biased towards termination in the orbitofrontal cortex, though the exact magnitude of the effect is more difficult to discern, owing to difficulties in reconstructing occipitotemporal-frontal connections using a data-driven methodology.

### Cross-species differences are robust against varying the orbitofrontal cortical region specification

A connection’s orbitofrontal bias index (OBI) may change depending on the specification of the orbitofrontal cortical and prefrontal cortical regions of interest. This is because OBI is calculated as a proportion of termination sites in the orbitofrontal cortex divided by the proportion of termination sites in the prefrontal cortex. As such, it is possible that a connection’s OBI might differ substantially depending on how the orbitofrontal and prefrontal cortical regions of index are specified.

To assess the extent to which our findings rely on the particularities of how we have defined the cortical masks, we performed an additional analysis in which we recalculated OBI values for each connection after redrawing the orbitofrontal cortical masks in the squirrel monkey and the lemur, the two species for which less cytoarchitectonic data are available to inform the specification of cortical masks. The new orbitofrontal masks (Supplemental Figure 3A) were drawn with particular care to exclude Brodmann Areas 9, 10, 24, 25, and 32, using the cytoarchitectonic maps reported by Rosabal (1967) for the squirrel monkey and Brodmann (1909) for the lemur.

Using these independently-specified masks, we observed OBI values as follows, presented as mean ± standard deviation. Anterior temporal-frontal connections in squirrel monkey: 2.01 ± 0.13; in lemur: 3.64 ± 0.87. Occipitotemporal-frontal connections in squirrel monkey: 1.80 ± 0.42; in lemur: 3.01 ± 1.21. The differences in OBI values resulting from this independent analysis are presented graphically in Supplemental Figure 3B. Nested ANOVA analyses showed significant differences between species on OBI values (anterior temporal-frontal connections – *F*: 30.30, *d*.*f*. = 2, p = 5.39 * 10^-6^; occipitotemporal-frontal connections – *F*: 16.08, d.f. = 2, *p* = 1.86 * 10^-4^). Post-hoc t-tests with the Bonferroni correction showed that with regional masks specified in this way, all species differed from one another (anterior temporal-frontal connections – macaque v. lemur: *p* = 2.40 * 10^-11^, squirrel monkey v. lemur: *p* = 1.90 * 10^-11^, macaque v. squirrel monkey: *p* = 8.03 * 10^-3^; occipitofrontal connections – macaque v. lemur: *p* = 1.00 * 10^-6^, squirrel monkey v. lemur: *p* = 8.59 * 10^-5^, macaque v. squirrel monkey: *p* = 8.85 * 10^-3^), with lemur OBI values being highest on both sets of connections. Thus, the finding holds that the ventral visual stream’s prefrontal terminations are most restricted to orbitofrontal cortex in the strepsirrhine; this finding does not seem to be particularly dependent on the way in which the orbitofrontal cortical masks are specified.

## Discussion

This work has used a data-driven analysis of tractography data in three primates to investigate the frontal projections of anterior temporal-frontal and occipitotemporal-frontal connections of the ventral visual pathway. In the absence of anatomical priors sufficient to specify exclusion masks and other criteria for atlas-based tractography in each of our three species, this technique has enabled us to compare the ventral visual pathway across species. We have shown that the frontal connections are far more widespread in the two anthropoids studied – the macaque and the squirrel monkey – than in the strepsirrhine primate – the lemur. Using the orbitofrontal cortex as a frontal reference region, we showed that the percentage of connections to this part of the frontal cortex is much larger in the strepsirrhine. These between-species effects were found both for anterior temporal-frontal and ventral occipital-temporal connections and were far greater in magnitude than either individual differences or differences due to parameterization of the analyses.

The primate temporal lobe has been argued to be a unique specialisation to that order (Bryant & Preuss, 2018). A large part of this cortical territory is devoted to the ventral visual stream, originating in the extrastriate cortex and extending anteriorly towards the temporal pole. In recent years, it has been increasingly appreciated that the ventral stream consists of a series of reciprocally connected regions, linked to other cortical and subcortical regions, acting as a recurrent network involved in the processing of object information (Kravitz et al., 2013). Complementing this understanding, the overall model of striate and extrastriate cortex representing visual features, and increasingly anterior temporal regions representing conjunctions of visual features and ultimately whole objects and even abstract representations, has proven robust (Braunsdorf et al., 2021; Murray et al., 2017).

Recent work has interpreted the ventral visual stream in terms of its role in visually based foraging in primates. The visual representations of the temporal cortex provide a sensory context that can be used to guide foraging decisions. It has been argued that the foraging niche of anthropoid primates is more complex than that of non-anthropoid primates, with anthropoids relying on more volatile sources of food and operating in a more complex landscape (Passingham & Wise, 2012). Indeed, anthropoid primates seem to have far more complex knowledge of their environment (Janmaat et al., 2006). Consistent with this, fossil evidence suggests that early anthropoids had smaller brains than modern anthropoids, yet still possessed the sharp visual acuity that underpins the complex foraging niche of modern anthropoids (Bush et al., 2004). This body of work is consistent with electrophysiological and anatomical studies. Eldridge et al. (2021) argue that many experimental results of neuronal activity in anthropoid temporal cortex, such as generalisation of items based on visual similarity and assessment of object properties based on visual information, are consistent with successful foraging behaviour in such an environment. Furthermore, Eldridge and colleagues (2021) note that the ventrolateral prefrontal cortex, as the other principal target of temporal visual cortical projections to the prefrontal cortex, is an anthropoid innovation absent in closely-related relatives, including strepsirrhines. Our results, showing that the anthropoid ventral visual stream involves relatively fewer projections to the orbitofrontal cortex than other prefrontal cortical regions, further develop the hypothesis that the ventral visual stream evolved in anthropoids to support sensory-driven foraging in complex environments.

More generally, our results demonstrate how data-driven methods for analysing complex neuroimaging data can be used successfully to test hypotheses in comparative neuroscience. Large-scale comparative anatomy is now possible through the increasing availability of high quality whole-brain imaging data from a range of species, including many primates (Assaf et al., 2020; Tendler et al., 2022). The availability of these data presents both opportunities and challenges, as many of the species from which data are now available have been studied only sparsely. The use of a data-driven approach, such as NMF, opens the possibility of interrogating these data without losing sight of anatomical rigour. Indeed, the inclusion of species that have been studied well, such as the macaque monkey, allows for a benchmarking or validation of methods. A similar approach has been taken before by Folloni and colleagues, showing how careful tractography data is able to reconstruct tracts previously identified using tracers in the macaque (Folloni et al., 2019), before expanding the approach to other species (Folloni et al., 2024).

The availability of multiple individuals per species also allowed us to assess the relative extent of within- and between-species variation in connectivity. Although the number of individuals per species in the current study is still quite low, this nevertheless allows a first quantification of the various contributions of variance in the data. To our knowledge, this has not been assessed in this manner before. We show that the between-species variance far outweighs that of individuals and of analysis parameterization. This result provides an important validation of earlier studies showing between-species differences in connectivity.

In sum, this work provides evidence of white matter reorganisation in the ventral visual stream in the anthropoid lineage. As the anthropoid primates evolved to occupy a niche as foragers capable of leveraging complex visual information, their ventral prefrontal cortex underwent extensive reorganisation. Our work confirmed that the ventral visual stream supporting this processing pathway also underwent changes in the anthropoid lineage. Furthermore, the data-driven techniques we have developed here hold the potential to address questions of white matter connectivity in additional, previously understudied species.

## Data availability

Analysis code is available under a CC BY-NC-SA 4.0 license at https://git.fmrib.ox.ac.uk/neuroecologylab/anthropoid-strepsirrhine-ventral-connectivity-public.

## Acknowledgements

JEH is funded by a Wellcome Trust Doctoral Studentship in Neuroscience (222368/Z/21/Z). The work of RBM was supported by the Biotechnology and Biological Sciences Research Council UK (BB/N019814/1) and the Medical Research Council UK (MR/Y010698/1). The Wellcome Centre for Integrative Neuroimaging is supported by core funding from the Wellcome Trust (203129/Z/16/Z). SW and SNS acknowledge funding from the European Research Council (ERC Consolidator 101000969 to SNS). The funders had no role in the design or writing of this report, nor in the collection, analysis, and interpretation of data. For the purpose of open access, the authors have applied a CC BY public copyright licence to any Author Accepted Manuscript version arising from this submission.

