## Supplemental Methods, Supplemental Figures 1-3, Supplemental Movie captions for "An anthropoid/strepsirrhine divergence in ventral visual stream connectivity"

#### Temporal and occipital region of interest specification

During component selection, regions of interest were drawn on each brain to screen for components that comprise anterior temporal-frontal and occipitotemporal-frontal connections. The regions of interest used in this screening process included prefrontal cortex and anterior temporal pole for anterior temporal-frontal connections, and prefrontal cortex and primary visual cortex for occipitotemporal-frontal connections. To identify whether this screening process might have missed anterior temporal-frontal connections with high signal intensity in other parts of temporal cortex and visual cortex, larger regions of interest were drawn to include more of the temporal and occipital cortex. Additional components identified in this screen were then subject to the same manual inspection for identification of putative anterior temporal-frontal and occipitotemporal-frontal connections.

#### Orbitofrontal region of interest in macaque

Because the macaque orbitofrontal cortex is smaller relative to the rest of the prefrontal cortex when compared against lemur and squirrel monkey, the macaque orbitofrontal region of interest was correspondingly smaller than in lemur or squirrel monkey. The macaque orbitofrontal cortex region of interest was 4.4% of the total prefrontal cortical region of interest, whereas the squirrel monkey and lemur orbitofrontal regions of interest were 23.3% and 19.9% of the prefrontal cortex, respectively.

It is possible that the disparity between species in the size of the orbitofrontal cortical regions of interest led to differences in reduced orbitofrontal bias index values in macaque that do not reflect a physiological difference in anterior temporal-frontal connections between species. As such, an additional analysis was performed using an expanded macaque orbitofrontal region of interest, comprising 23.0% of the prefrontal cortical area. Supplemental Figure 2 presents a visual comparison of the standard and expanded orbitofrontal regions of interest in macaque.

### Supplemental Figures

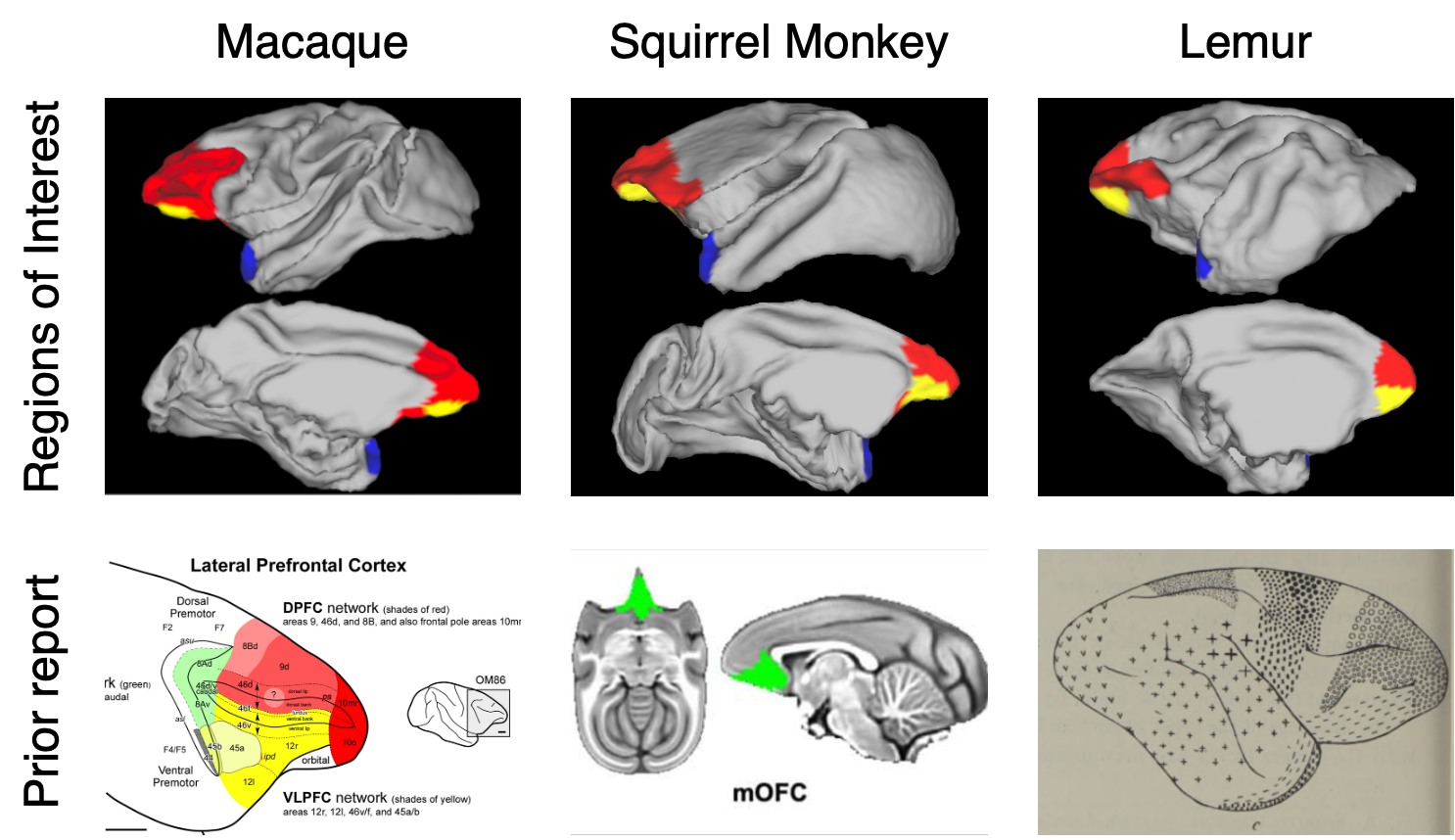

**Supplemental Figure 1. Comparison between regions of interest and prior literature.** Top row: Left hemisphere regions of interest for macaque, squirrel monkey, and lemur. Bottom row: representative images from prior publications used in specifying regions of interest. Macaque image from Saleem et al. (2014). Squirrel monkey image from Deshpande and Kohut (2023). Lemur image from Mott & Kelley (1908). Yellow = orbitofrontal cortex. Red = prefrontal cortex. Blue = anterior temporal pole. mOFC = medial orbitofrontal cortex.

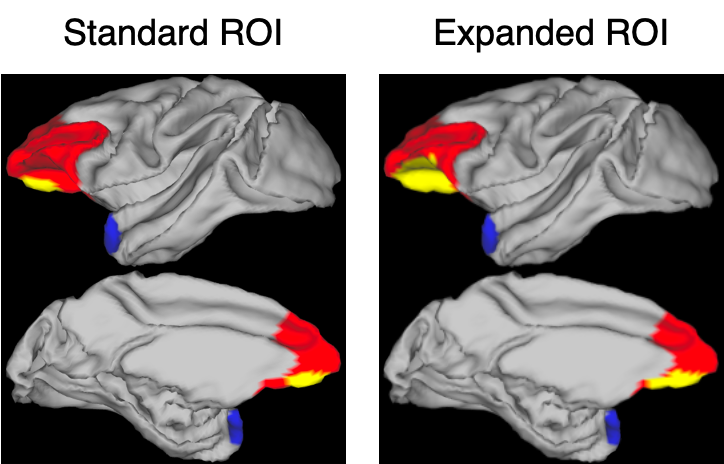

**Supplemental Figure 2. Comparison between standard and expanded orbitofrontal regions of interest in macaque.** Surface projections showing regions of interest (ROIs) used to screen NMF components. All images show the lateral surface (above) and medial wall (below) of the left hemisphere. The standard orbitofrontal region of interest (yellow) comprised 4.4% of prefrontal cortex (red), while the expanded orbitofrontal region of interest comprised 23.0% of prefrontal cortex. Blue indicates the anterior temporal pole region of interest, which was unchanged for this analysis.

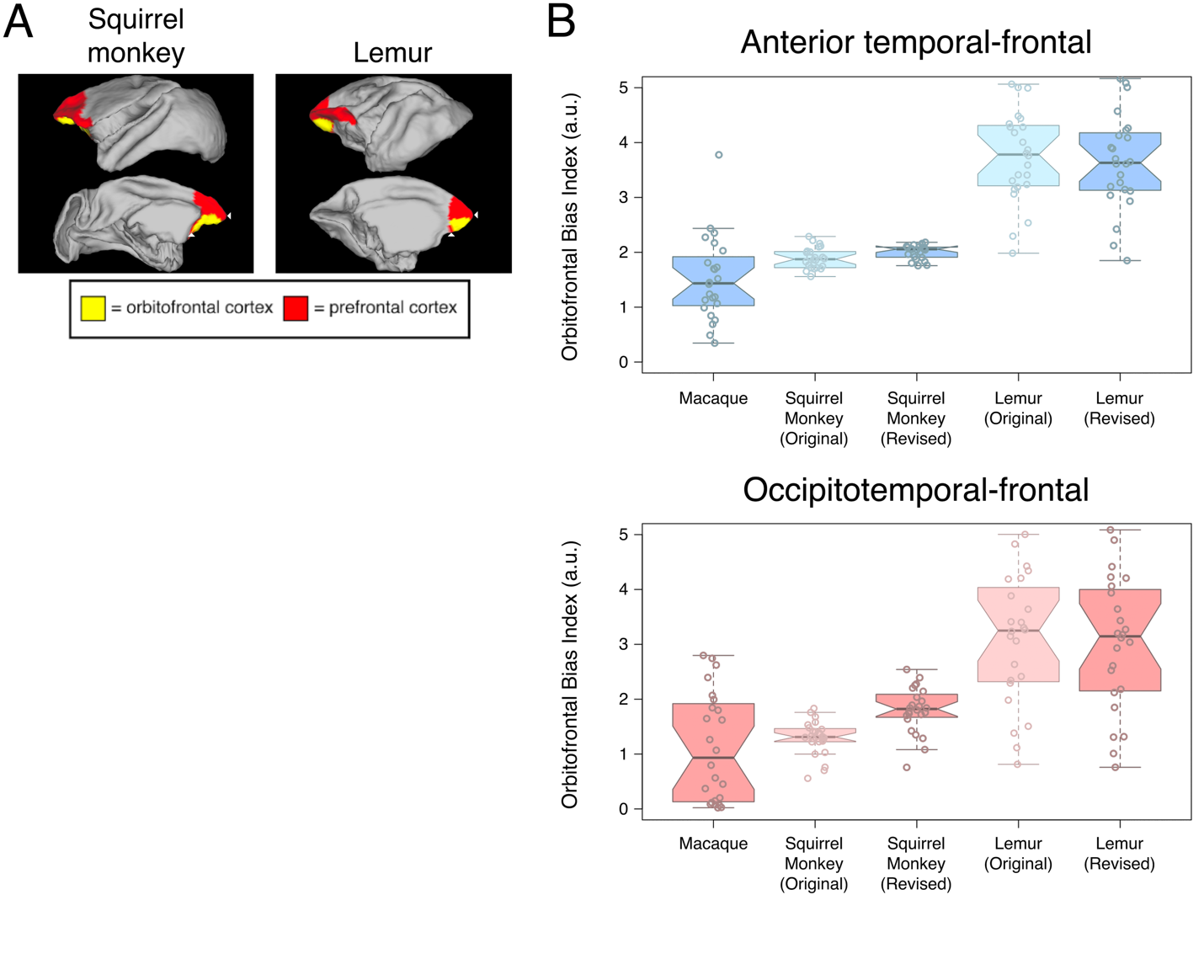

**Supplemental Figure 3. Alternative orbitofrontal cortical region specification.** **A.** Alternative orbitofrontal cortical regions of interest in squirrel monkey and lemur, specified according to Rosabal (1967) and Brodmann (1909). White arrow heads indicate notable differences from the original masks at the frontal pole and posterior section of prefrontal cortex. **B.** Box-and-whisker plots show the distributions of orbitofrontal bias index values. Lighter, less-saturated plots indicate values attained using the original orbitofrontal cortical masks, presented here for visual comparison. Statistical analysis included only the squirrel monkey and lemur values which have been labelled “revised”. Notches indicate the 95% confidence interval of the median (median ± 1.57 * IQR/n^0.5^).

**Supplemental Movie 1. Anterior-to-posterior coronal sections in the macaque brain.** Presented in radiological view (i.e., flipped on the horizontal axis). Blue indicates occipitotemporal-frontal connections, orange indicates anterior temporal-frontal connections.

**Supplemental Movie 2. Dorsal-to-ventral axial sections in the macaque brain.** Presented in radiological view (i.e., flipped on the horizontal axis). Blue indicates occipitotemporal-frontal connections, orange indicates anterior temporal-frontal connections.

**Supplemental Movie 3. Left-to-right sagittal sections in the macaque brain.** Blue indicates occipitotemporal-frontal connections, orange indicates anterior temporal-frontal connections.

**Supplemental Movie 4. Anterior-to-posterior coronal sections in the squirrel monkey brain.** Presented in radiological view (i.e., flipped on the horizontal axis). Blue indicates occipitotemporal-frontal connections, orange indicates anterior temporal-frontal connections.

**Supplemental Movie 5. Dorsal-to-ventral axial sections in the squirrel monkey brain.** Presented in radiological view (i.e., flipped on the horizontal axis). Blue indicates occipitotemporal-frontal connections, orange indicates anterior temporal-frontal connections.

**Supplemental Movie 6. Left-to-right sagittal sections in the squirrel monkey brain.** Blue indicates occipitotemporal-frontal connections, orange indicates anterior temporal-frontal connections.

**Supplemental Movie 7. Anterior-to-posterior coronal sections in the lemur brain.** Presented in radiological view (i.e., flipped on the horizontal axis). Blue indicates occipitotemporal-frontal connections, orange indicates anterior temporal-frontal connections.

**Supplemental Movie 8. Dorsal-to-ventral axial sections in the lemur brain.** Presented in radiological view (i.e., flipped on the horizontal axis). Blue indicates occipitotemporal-frontal connections, orange indicates anterior temporal-frontal connections.

**Supplemental Movie 9. Left-to-right sagittal sections in the lemur brain.** Blue indicates occipitotemporal-frontal connections, orange indicates anterior temporal-frontal connections.
